# Autometa: Automated extraction of microbial genomes from individual shotgun metagenomes

**DOI:** 10.1101/251462

**Authors:** Ian J. Miller, Evan R. Rees, Jennifer Ross, Izaak Miller, Jared Baxa, Juan Lopera, Robert L. Kerby, Federico E. Rey, Jason C. Kwan

## Abstract

**Motivation:** Shotgun metagenomics is a powerful, high-resolution technique enabling the study of microbial communities *in situ.* However, species-level resolution is only achieved after a process of “binning” where contigs predicted to originate from the same genome are clustered. Such culture-independent sequencing frequently unearths novel microbes, and so various methods have been devised for reference-free binning. Existing methods, however, suffer from: (1) reliance on human pattern recognition, which is inherently unscalable; (2) requirement for multiple co-assembled metagenomes, which degrades assembly quality due to strain variance; and (3) assumption of prior host genome removal not feasible for non-model hosts. We therefore devised a fully-automated pipeline, termed “Autometa,” to address these issues. Results: Autometa implements a method for taxonomic partitioning of contigs based on predicted protein homology, and this was shown to vastly improve binning in host-associated and complex metagenomes. Autometa’s method of automated clustering, based on Barnes-Hut Stochastic Neighbor Embedding (BH-tSNE) and DBSCAN, was shown to be highly scalable, outperforming other binning pipelines in complex simulated datasets.

**Availability and implementation:** Autometa is freely available at https://bitbucket.org/jasonckwan/autometa and as a docker image at https://hub.docker.com/r/jasonkwan/autometa under the GNU Affero General Public License 3 (AGPL 3).

**Supplementary information:** Supplementary data are available attached to this article at https://biorxiv.org

## 1 Introduction

Microbes are known to associate with almost all organisms on Earth, including humans, where they are thought to have tremendous impact in health, disease, and agriculture (Dubilier *et al.*, 2015; Alivisatos *et al.*, 2015; Buick, 2008). However, it has long been known that only a minute fraction of environmental microbes are readily cultured in the laboratory (Staley and Konopka, 1985). Thus, the vast majority of the microbial tree of life is as yet only accessible through culture-independent sequencing (“metagenomics”). Early metagenomic studies focused on phylogenetic profiles of communities by examining the relative abundance of individual bacterial species within different environments (quantified through 16S rRNA gene sequencing), but offered limited information about the functional contribution and organism-level interactions that shape these environments (Escobar-Zepeda *et al.*, 2015). Whole genome “shotgun” sequencing is able to overcome some of the challenges faced by high-throughput 16S rRNA amplicon sequencing, such as the issue with non-canonical ribosomal RNA genes that are undetectable through standard primers (Brown *et al.*, 2015) and the inherent low-resolution nature of single gene studies. However, the task of sorting metagenomic contigs into clusters representing individual genomes (“binning”) is a challenging computational problem and an active area of research (Sedlar *et al.*, 2017; Sangwan *et al.*, 2016). Binning is a necessary step towards understanding the metabolic and functional contributions of individual microorganisms to metabolic capabilities of the community as a whole. In other words, genome-level resolution of metagenomes allows researchers to move beyond the interpretation of metabolic function in aggregate to understanding the role of individual organisms within a complex system *in situ*.

Given that most environments are predominantly composed of uncharacterized microorganisms, different approaches have been taken to achieve reference-free binning. For instance, nucleotide composition has been used to group contigs with emergent self-organizing maps (ESOM) (Dick *et al.*, 2009) or Barnes-Hut stochastic Neighbor Embedding (BH-tSNE) (Laczny *et al.*, 2014, 2015). These approaches reduce variation in *k*-mer frequencies to two dimensions, enabling the visualization of highly dimensional data and allowing human-driven clustering. Other efforts have focused on leveraging information from multiple samples, with the assumption that contigs in shared genomes will show a distinct co-variance in coverage. Both manual (Nielsen *et al.*, 2014; Albertsen *et al.*, 2013) as well as automatic pipelines (Wu *et al.*, 2014; Alneberg *et al.*, 2014) have used this approach. However, there are a number of disadvantages to this methodology. Many multi-sample protocols require assembly of reads from all samples (referred to as “co-assembly”), increasing computational requirements and potentially degrading assembly quality when shared genomes are not clonal. This issue is known as “microdiversity” - a problem acknowledged by (Albertsen *et al.*, 2013) and recently demonstrated elsewhere (Olm *et al.*, 2017). By pooling samples for co-assembly, users can also exacerbate the effect of population summing, whereby a genome assembly represents broadly aggregated consensus sequences instead of the genome of a single strain, organism, or population taken from one sample (Sangwan *et al.*, 2016; Miller *et al.*, 2017; Olm *et al.*, 2017). Such aggregation can mask the presence of pan-genome sequences (Medini *et al.*, 2005) found only in individual strains or samples, which has important clinical and biotechnological implications when considering mobile elements that confer antibiotic resistance (Tettelin *et al.*, 2008; Moss *et al.*, 2017) or biosynthetic gene clusters acquired through horizontal transmission. There are further situations where the underlying variability or overlap of the system is unknown, and there is a desire to extract information from a small number of pilot datasets. Additionally, multi-sample comparisons, which by nature incur higher sequencing costs, do not necessarily aid in binning of genomes unique to one sample (Miller, Weyna, *et al.*, 2016).

To date, our efforts to sequence the genomes of marine invertebrate symbionts that make bioactive small molecules have relied upon semi-manual binning techniques (Miller, Vanee, *et al.*, 2016; Miller, Weyna, *et al.*, 2016). However, marine sponge microbiomes, which offer a wealth of biotechnological potential (Lackner *et al.*, 2017), can contain hundreds of microbial species, occupying up to 40% of the sponge’s tissue volume (Hentschel *et al.*, 2012; Taylor *et al.*, 2007). As these systems were beyond the limit of reasonable manual processing, and due to the poor performance of existing automatic binning pipelines for such host-associated metagenomes, we were motivated to develop an automated and scalable binning algorithm, which we call “Autometa”. This method carries out clustering on a simplified subset of contigs (those taxonomically classified as either Bacteria or Archaea), in order to maximize scaling according to metagenomic complexity from individual metagenome assemblies. The initial clusters serve as the training set for subsequent classification by a supervised machine learning algorithm. We evaluated Autometa using a number of simulated and synthetic metagenomes, where performance could be assessed with reference to the known component genomes, as well as a real host-associated metagenome we previously examined by semi-manual binning (Miller, Vanee, *et al.*, 2016; Miller, Weyna, *et al.*, 2016). We found that Autometa performed comparably or outperformed MaxBin (Wu *et al.*, 2014), MetaBAT (Kang *et al.*, 2015), MyCC (Lin and Liao, 2016), and BusyBee Web (Laczny *et al.*, 2017), especially in cases with higher metagenome complexity and in a host-associated dataset. We further found that contig-level taxonomic classification using lowest common ancestor (LCA) analysis was able to improve Autometa’s performance as well as the binning performance of other pipelines.

## 2 Methods

Autometa bins microbial genomes *de novo* from single shotgun metagenomes using sequence homology, coverage, and nucleotide composition to distinguish between contigs. The task is guided by the presence of marker genes, previously identified in Bacteria and Archaea (Rinke *et al.*, 2013) and known to occur as single copies in microbial genomes. The presence of marker genes can be used to estimate the genome completeness of bins, as well as the level of contamination, as each marker should only be detected once per bin. The microbes found in environmental metagenomes can be highly divergent from all previously sequenced organisms, and those that associate with eukaryotic hosts often undergo a process of genome degradation and reduction, where functions essential to independent life can be lost (McCutcheon and Moran, 2012; Bennett and Moran, 2015). For instance, we recently identified a genome-reduced bacterium that was so divergent from known sequences that only ~20% of genes had hits in the NCBI NR database, and only ~20% of the expected bacterial single-copy markers could be detected (Miller, Weyna, *et al.*, 2016). We therefore do not assume bins should be close to 100% complete or use single copy markers to pre-calculate the number of bins, as in MaxBin (Wu *et al.*, 2014). The overall process employed in Autometa comprises three broad stages (**Figure 1**):

1. Separate contigs into kingdom bins based on sequence homology.
2. Iteratively cluster kingdom-specific contigs.
3. Classify unclustered contigs to bins via supervised machine learning.

**Figure 1.**
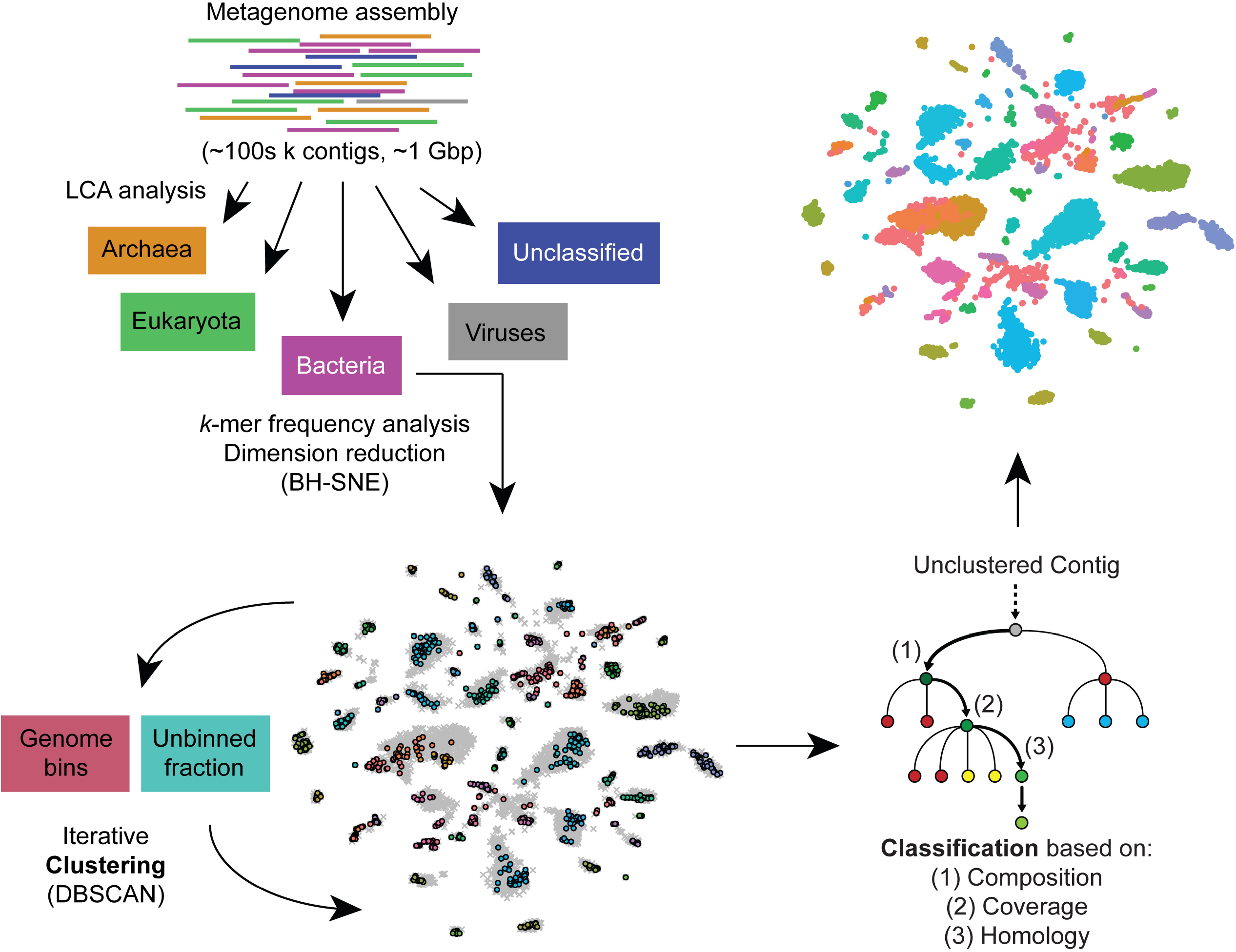
Autometa binning workflow. Autometa separates contigs from a *de novo* metagenome assembly into kingdom-level bins based on sequence homology, iteratively clusters kingdom-specific (Bacterial or Archaeal) contigs, and then (optionally) classifies any remaining unclustered contigs to bins using a decision tree classifier.

### 2.1 Separate contigs into kingdom bins

A broad separation of contigs into kingdom bins allows the removal of host-derived or other eukaryotic contamination (even if the host genome is not represented in reference databases), as well as separation of contigs derived from Bacteria and Archaea, simplifying subsequent deconvolution. Genes are identified in all contigs longer than a specified length cutoff (default is 10,000 bp, all datasets tested here were based on a 3,000 bp cutoff) with Prodigal (Hyatt *et al.*, 2010). Translated coding sequences are then queried against the NCBI NR database using the accelerated BLAST implementation Diamond (Buchfink *et al.*, 2015). The LCA of the hits with bitscore within 10% of the top hit is used to assign a taxonomy ID to each predicted protein according the the NCBI taxonomy database (https://www.ncbi.nlm.nih.gov/taxonomy). To reduce the influence of horizontally transferred genes, contig-level taxonomy is assigned by a modified majority vote of the component predicted coding sequences. Classifications are considered in order of decreasing specificity (species, then genus, family, order, class, phylum and kingdom), and accepted when a majority (≥50%) classification is reached, provided that the majority of proteins classified with lower specificity are ancestors of this classification. If an answer cannot be reached by this process, the lowest common ancestor of all proteins within a contig is used as the contig classification. Because eukaryotic genomes have low coding density, this system might conceivably lead to incorrect assignment of eukaryotic contigs as bacterial/archaeal in the case of interkingdom HGT. While a filter for coding density might distinguish most bacterial contigs from eukaryotic ones, employing the wrong cutoff would exclude low-density symbiont genomes at early points in genome reduction (McCutcheon and Moran, 2012). Most identified bacterial to eukaryotic HGT events are from organelles (K. B. Sieber *et al.*, 2017), and therefore we anticipate that in these cases the closest BLAST hits will be other organelle genes, tied to the host taxonomy. Additionally, the use of prokaryotic gene-finding algorithm Prodigal is expected to yield multiple ORFs for each eukaryotic gene, corresponding to each exon, thus potentially weighting eukaryotic classifications over prokaryotic ones. Very divergent prokaryotic genomes can contain contigs with varying classification even at the phylum level (Miller, Weyna, *et al.*, 2016), and therefore taxonomic classification is used cautiously in subsequent operations (see below). At this stage, contigs are separated into bins classified according to different kingdoms, and contigs classified as Bacteria and/or Archaea are progressed to the next step.

### 2.2 Clustering kingdom-specific contigs

It has been shown that *k*-mer frequency patterns differ between bacterial species/strains (Teeling *et al.*, 2004; Dick *et al.*, 2009), and that visualization of *k*-mer frequency data after dimension reduction with Barnes-Hut Stochastic Neighbor Embedding (BH-tSNE) (Van Der Maaten, 2014) effectively aids manual deconvolution of metagenomic contigs (Laczny *et al*., 2014, 2015). However, the feasibility and throughput of visual (manual) binning using BH-tSNE quickly degrades with increasing metagenome complexity. Autometa counts 5-mer frequencies in contigs, normalizes and reduces the raw dimensions to 50 with principal component analysis (PCA) as previously described (Laczny *et al.*, 2014), before dimension reduction with BH-tSNE.

In BH-tSNE, the parameter of “perplexity” can be conceptualized as the effective number of neighbors considered when the algorithm embeds local structure. In previous work (Laczny *et al.*, 2014), a perplexity value of 30 has been used, so we sought to determine if this was a reasonable value to use in all cases, or whether the parameter should be optimized for different datasets. It has been suggested (Cao and Wang, 2017) that a factor referred to as pseudo Bayesian Information Criteria (pBIC, or S) might be used to determine the optimum perplexity value (judged by human machine learning experts), where the optimum perplexity gives the minimum value of S. In simulated and synthetic metagenomes (see below), we found that minimum S values scale with the number of contigs (Figures S1–S6). To determine whether this would be a valid method for optimizing perplexity, we devised an objective measure of separation based on alignments of metagenomic contigs to the input genomes. For a particular perplexity, we construct groups of points in BH-tSNE space based on their assigned genomes (discounting contigs that are misassembled or unalignable). Within the groups, we discard outliers whose distance away from the group’s centroid is greater than the third quartile of distances plus the interquartile range multiplied by 1.5. From the remaining points, a convex hull is constructed, and we determine both the total area of the hull, *t*, and the area that is not overlapped by any other genome convex hull, *u*. The ‘non-overlapping fraction’, v, of the coordinate set for a given perplexity is given by equation 1, where the optimum perplexity should yield the maximum value of v, representing the greatest separation between genomes in BH-tSNE space. We plotted *v* against perplexity for all datasets where a ground truth was known (see below, Figures S7–S12). Importantly, peak values of *v* did not occur at perplexities close to those giving minimum values for S, meaning that *S* is a poor predictor for *v*. A degree of variability in adjacent values of perplexity was observed, due to the stochastic nature of BH-tSNE, but the peak value of *v* generally occurred between perplexities of 20 and 70, regardless of dataset size. As a relationship between dataset size and optimal perplexity was not found, and a method for optimizing perplexity in the absence of ground truth information was not apparent, we use a default value of 30 in the Autometa pipeline.

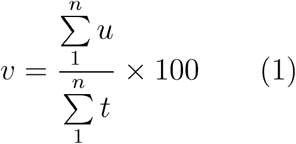

Clustering is achieved with the DBSCAN algorithm (Ester *et al.*, 1996), which clusters based on local density and is able to exclude outliers. In other words, it does not force all contigs into a bin, minimizing the potential for overfitting. DBSCAN has been previously implemented to cluster the output of dimension reduction of pentanucleotide frequencies via BH-tSNE (Heintz-Buschart *et al.*, 2016). Here, as input to the DBSCAN algorithm, we use the two dimensions produced by BH-tSNE as well as contig coverage. The eps parameter for DBSCAN controls the size of the local neighborhood around points that is explored during clustering, and we cycle through ascending values of eps from 0.3, increasing by 0.1 until only one group is obtained. For each of these iterations, Autometa assesses clusters by examining both their completeness (number of expected single copy markers) and purity (number of single copy markers that are unique in the cluster). The eps value selected is the one that gives the highest median completeness of bins that are above 20% complete and 90% pure, and the resulting bins that pass these criteria are kept. This method has the advantage that it assesses clustering in a biologically relevant manner, in contrast to internal clustering validation functions (Liu *et al.*, 2010), and it balances recall (completeness) and precision (purity) of the resulting bins. The clusters that do not meet these criteria and contigs in the “unclustered” bin are then subjected to another round of DBSCAN, again maximizing for median completeness of clusters over 20% complete and 90% pure. This process is iterated until no more clusters meeting the completeness and purity criteria can be obtained. In our investigations of the effects of perplexity on *v* (Figures S7–S12), we found that peak values of *v* decreased with increasing dataset size, illustrating that BH-tSNE is not able to avoid spatial overlap in complex datasets. In these cases it is expected that further fractionation of the data based on orthogonal properties will improve clustering quality. Therefore, we allow the unclustered fraction in each iteration to be optionally further divided into taxonomic groups in ascending order of specificity (phylum, then class, order, family, genus and species). After each split the iterative DBSCAN algorithm described above is repeated, and if unclustered sequences result, they are pooled for clustering at the next specific taxonomic level. This process allows the deconvolution of taxonomically-distinct genomes that exhibit similar *k*-mer frequency and coverage, and starts at the non-specific end of the taxonomic spectrum (i.e., phylum before class) to yield well-separated clusters and to maximize the chance of clustering divergent genomes that exhibit uncertain taxonomic classification (see above).

### 2.3 Classifying remaining contigs by supervised machine learning

Following initial clustering of contigs into bins, all remaining (i.e., unclustered) contigs are further recruited to these cores using a supervised decision tree classifier approach. The classifier is trained with features of single copy gene marker-containing contigs belonging to the initial clusters, using 5-mer frequencies reduced to 50 dimensions via PCA, as well as sequence coverage, and (optionally) taxonomic information encoded as a binary indicator matrix. The confidence of each of the classifier’s predictions is measured using Jackknife cross validation whereby the classifier is iteratively re-trained with a random subset (50%) of the training data (Chevrette *et al.*, 2017). By default, a prediction will only be accepted if this metric reports 100% confidence (e.g., 10/10 consistent classifications when trained with 10 random subsamples of the training data) and the prediction does not add any marker contamination to the predicted bin. After each full round of predictions, any marker-containing contig that is confidently classified to pre-existing clusters is added to the training data for subsequent rounds of classification, until no further marker-containing contigs are confidently classified. This approach is similar to the ‘bootstrapping’ of supervised machine learning in BusyBee web using the result of unsupervised clustering, except that it includes features beyond nucleotide composition, such as sequence coverage and taxonomic information in the prediction process and uses jackknife cross validation (**Figure S13**) to assess the confidence of each prediction.

### 2.4 Benchmarking

#### 2.4.1 Benchmarking datasets

Simulated metagenomes of increasing complexity (see **Table 1**) were created by picking random genomes out of the bacterial genome assemblies held in the NCBI database. Illumina reads (2 × 125 bp) were simulated using ART (Huang *et al.*, 2012) and assembled with metaSPAdes (v3.9.0) (Nurk *et al.*, 2017). Datasets were simulated to represent each component genome with equal coverage in order to stress test discrimination performance based on nucleotide composition. A synthetic metagenome (“Mix-51”) was made by mixing together 51 bacteria isolated from the human gut, before extracting DNA as previously described (McNulty *et al.*, 2011). The resulting DNA pellet was dissolved in TE buffer (10 mM Tris-HCl pH 8.0, 1 mM EDTA) then column-purified using the Nucleospin Gel and PCR Clean-up kit (Macherey-Nagel Inc, Bethlehem, PA). DNA concentration was measured using the Qubit BR dsDNA assay (Invitrogen, Eugene, OR). Sequencing of this DNA was carried out on an Illumina HiSeq 2500, in a 2 × 125 bp run. Adapters were trimmed from the resulting reads using Trimmomatic (Bolger *et al.*, 2014), before being assembled with metaSPAdes (Nurk *et al.*, 2017). The raw reads of Mix-51 are accessible through the Sequence Read Archive (SRA), under accession number **SRR5679054**. Further information on the datasets, including details of component genomes, can be found in **Table S1**.

**Table 1.**
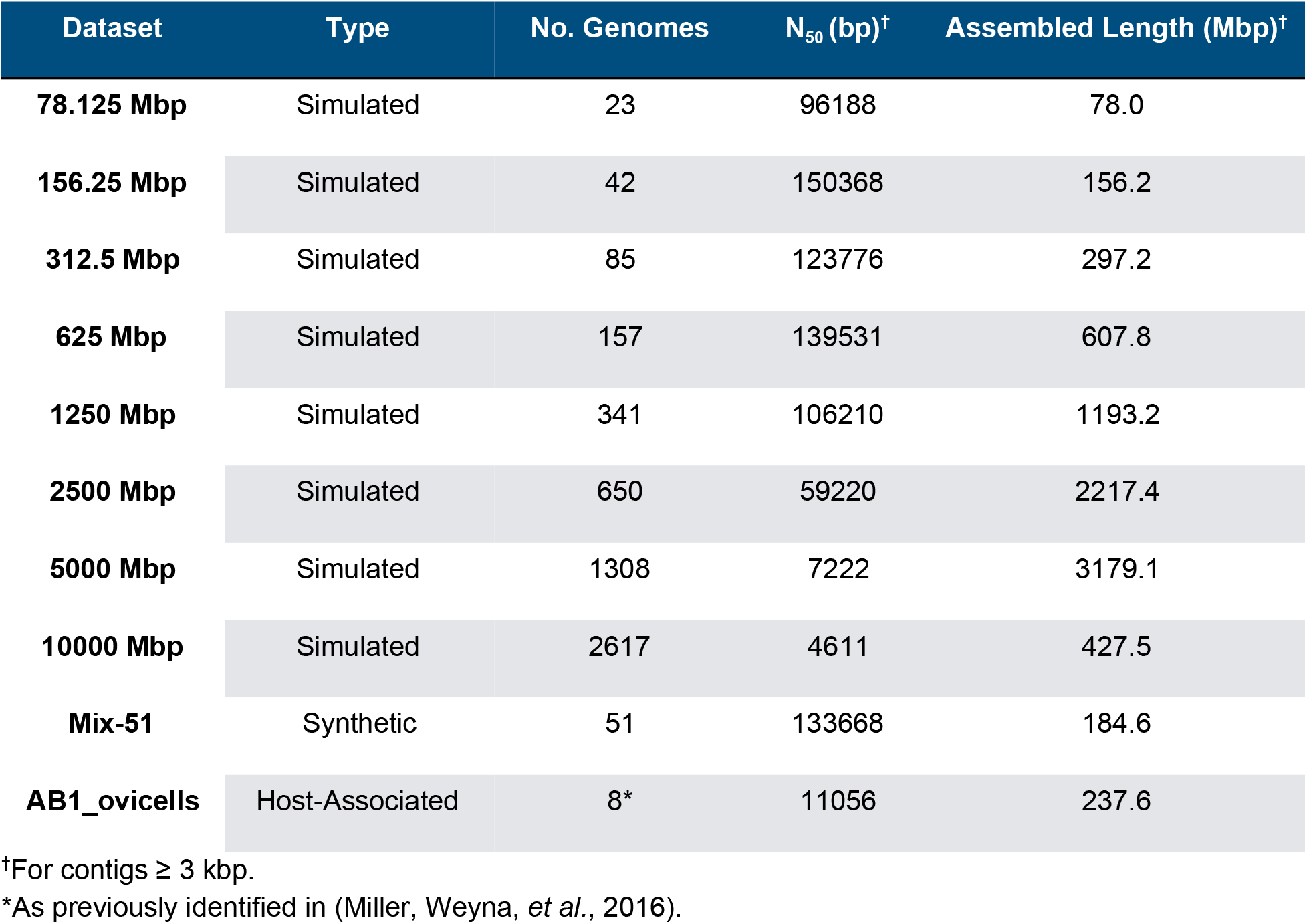
Datasets used in this study.

We also included sample AB1_ovicells in benchmarks, which we previously examined semi-manually (Miller, Vanee, *et al.*, 2016; Miller, Weyna, *et al.*, 2016). This is a metagenome associated with a marine bryozoan, containing the uncultured bryostatin-producing symbiont, *“Candidatus* Endobugula sertula” along with several divergent bacteria and several genomes that are very similar in GC content and/or coverage. The same assembly used previously (Miller, Vanee, *et al.*, 2016; Miller, Weyna, *et al.*, 2016) was assessed as a point of comparison to manual binning efforts. Previously published raw reads and annotated assemblies associated with sample AB1_ovicells are available through NCBI (BioProject **PRJNA322176**). All datasets were tested using commit version 9592e35 and run on a linux server (Dell Poweredge T430 with two Intel Xeon E5-2650 v3 2.3 GHz CPUs, 128 GB of RAM and 1.7 TB of disk space).

#### 2.4.2 Choice of comparison pipelines

To enable an apposite comparison of Autometa’s performance with existing pipelines, we excluded pipelines with different aims, such as those designed to pre-cluster raw sequence reads or those that required multiple metagenomic datasets. We also excluded pipelines that required manual interpretation of visualizations, on the grounds that these did not include an automated clustering step. This rationale led us to focus on four pipelines for comparison: MaxBin (Wu *et al.*, 2014), MetaBAT (Kang *et al.*, 2015) MyCC (Lin and Liao, 2016) and BusyBee Web. (Laczny *et al.*, 2017).

#### 2.4.3 Evaluation metrics

In the case of the simulated metagenomes and Mix-51, contigs were assigned to reference genomes with metaQUAST (Mikheenko *et al.*, 2016). Precision and recall were then calculated as described previously (Lin and Liao, 2016) according to **equations 2** and **3** where we consider the binning of *N* genomes into *M* clusters and *S_ij_* is the combined length of contigs in cluster *i* which belong to reference genome *j*. Precision is a property of clusters, described as the length fraction of a cluster taken up by contigs belonging to the genome accounting for the largest fraction of the cluster (*max_j_*). Recall is a property of reference genomes, described as the length fraction of the genome assigned to the cluster with the largest fraction of that genome (*max_j_*). For the purposes of these calculations, contigs labelled as “misassembled” by MetaQUAST were excluded. The F1 score is the harmonic mean of precision and recall (**equation 4**).

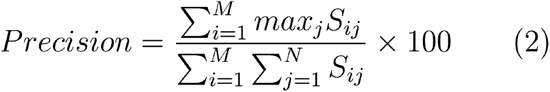

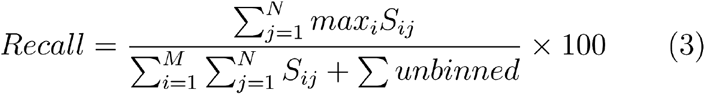

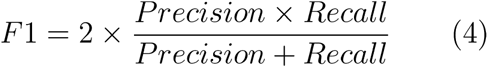

## 3 Results

### 3.1 Performance in a host-associated metagenome

Compared to our previous semi-manual binning efforts of the AB1_ovicells sample, all four tested programs produced a greater number of bins. Autometa produced 22 genome bins (**Table 2**) compared to the eight we identified by our earlier, semi-manual approach (**Figure 2** and **Table 1**). However, when comparing the performance of these programs to the composition of manually classified sequence, the bin-level performance was more variable. For instance, each program performed differently when compared to our semi-manual classification of “Ca. Endobugula sertula” contigs. We defined the original AB1_ovicells *“Ca.* E. sertula” assembly to include 3.32 Mbp in 117 contigs (Miller, Vanee, *et al.*, 2016). The cluster statistics of the “Ca. E. sertula” bin as identified by the four automated different programs are detailed in **Table S2**. Autometa produced the genome bin most consistent with the semi-manual binning (recovering 92/117 contig (93.3% of length) derived from semi-manual binning). MaxBin had the second highest recovery of the original *“Ca.* E. sertula” assembly, at a 91.9% recovery rate. Autometa and MaxBin were tied for the highest apparent completeness for this cluster, at 96.2%. The “Ca. E. sertula” cluster identified by MyCC had a slightly higher purity (98.2% compared to 96.6%, according to CheckM results), but with the lowest completeness (71.6%). Interestingly, both MyCC and Autometa identified a shared set of 44 contigs, within the *“Ca.* E. sertula” bin, that we had previously left unclassified through our semi-manual efforts. The nucleotide composition of these contigs was consistent with contigs we identified as belong to *“Ca.* E. sertula” (**Figure S14**), but had lower sequence coverage on average (**Figure S15**). However, 12 of these 44 contigs (27%) identified by Autometa and MyCC were assigned the order level taxonomy of “Oceanospirillales,” which suggests contamination from another gammaproteobacteria genome bin that we previously identified as an *Endozoicomonas* sp. (**Figure S16**).

**Figure 2.**
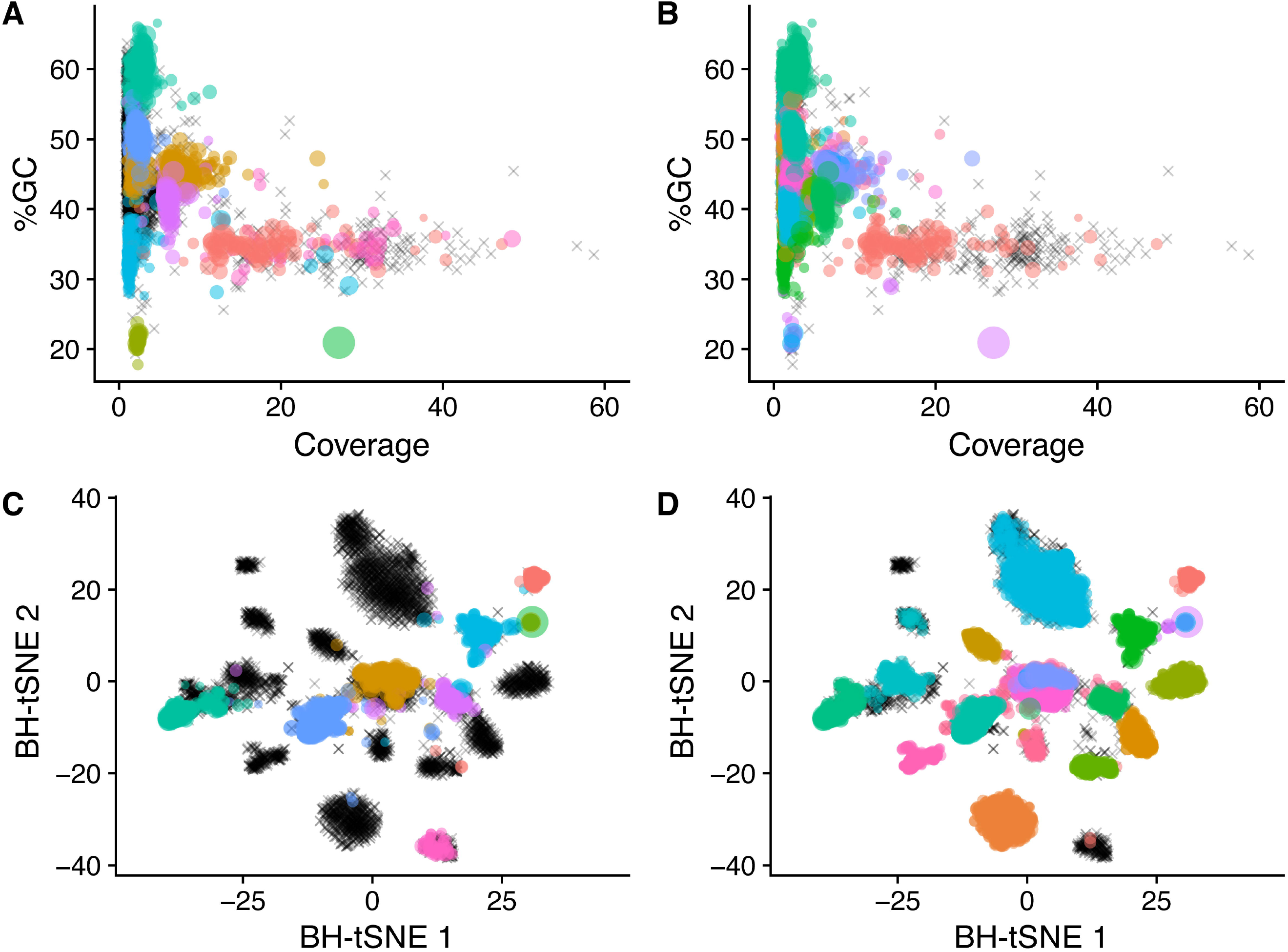
Visualization of genome bins from a host-associated metagenome derived from semi-manual binning vs. automated binning. Points represent contigs and are colored based on their assigned bin and scaled by length; unclustered contigs are represented by black crosses.

**Table 2.** Effect of taxonomic partitioning on binning performance of AB1_ovicells.

| Algorithm | AB1_ovicells<br>with taxonomy filtering |  |  | AB1_ovicells<br>without taxonomy filtering |  |  |
| --- | --- | --- | --- | --- | --- | --- |
|  | No. bins | Median<br>Completeness | Median<br>Purity | No. bins | Median<br>Completeness | Median<br>Purity |
| Autometa | 22 | 34.0 | 99.6 | 20 | 40.9 | 99.1 |
| MyCC | 25 | 31.0 | 98.8 | 22 | 41.1 | 96.3 |
| MaxBin | 13 | 74.3 | 88.1 | 27 | 23.7 | 95.6 |
| MetaBAT | 25 | 23.6 | 99.9 | 40 | 13.2 | 100.0 |

The aggregate binning results for Autometa and MyCC for the AB1_ovicells sample appear comparable in the number of bins recovered, along with median purity and completeness metrics, with an apparent tradeoff between purity (higher with Autometa) and completeness (higher with MyCC). However, it is worth noting that CheckM does not systematically consider contamination from host Eukaryotic sequences in its reported contamination statistics and much of the sequence clustered by MyCC, MaxBin, and MetaBAT appears contaminated with Eukaryotic sequence (**Table S3**). In fact, at least two MyCC clusters (Cluster.8 and Cluster.2, **Table S4**) appear heavily contaminated by host bryozoan sequence, though CheckM reports their purity as 87.5% and 77.3% and marker lineage as Archaea (**Table S4**). These clusters represent 136.3 Mbp and 31.7 Mbp, in 16,825 and 2,811 contigs, respectively (**Table S4**). Analysis with Autometa’s LCA workflow suggests only 3.5 Mbp (2.5%), and 1.8 Mbp (5.8%), respectively, of these MyCC clusters are represented by microbial sequence (**Table S5**).

Thus, to test the effect of taxonomic filtering on the binning performance of other pipelines, we repeated runs with MaxBin, MetaBAT, and MyCC on just contigs that Autometa’s LCA workflow identified as bacterial. This taxonomic filtering step resulted in decreased bin fragmentation for MaxBin and MetaBAT. In addition to preventing these putative host sequences from populating MyCC bins, taxonomic filtering improved the median cluster statistics for MetaBAT and MaxBin. Without taxonomic filtering, MetaBAT identified 40 genome bins with an median completeness of 13.2, compared to the 25 bins identified with an median completeness of 23.6 with taxonomy-filtered contigs (**Table 2**). Taxonomic filtering also resulted in a consolidation of bins produced by MaxBin, with a concomitant increase in completeness (74.3% with and 23.7% without taxonomic filtering).

### 3.2 Performance in a synthetic metagenome with highly similar strains

All tested algorithms were challenged by the high strain overlap of the synthetic Mix-51 community (**Figure 3, Table S6**). Autometa produced 57 genome bins with an average completeness of 72.3% and an average purity of 99.6%, based on CheckM analysis, where the completeness percentage here is affected by the percentage of the genome that was effectively assembled (Parks *et al.*, 2015). For 33 of the 51 bacteria in Mix-51, Autometa produced the highest quality bins based on F1 value, with a median F1 value of 0.935, compared to median values of 0.571, 0.763, 0.694, and 0.324 of MyCC, MaxBin, and MetaBAT, and BusyBee Web respectively (**Table S6**). Of the closely related *Bacteroides* species, Autometa best resolved the genomes of *B. stercoris, B. uniformis, B. cellulosilyticus*, and *B. intestinalis.* However, all four algorithms struggled to classify the closely related *B. thetaiotaomicron* strains, which also exhibited some of the lowest fractions of assembly (**Table S6**). For instance metaSPAdes was only able to recover 59.9% of the Bacteroides thetaiotaomicron 3731 genome, based on alignments of the *de novo* assembly to known reference genomes using metaQUAST (Mikheenko *et al.*, 2016). In terms of F1 values, MaxBin consistently outperformed all other algorithms in resolving these three strains.

**Figure 3.**
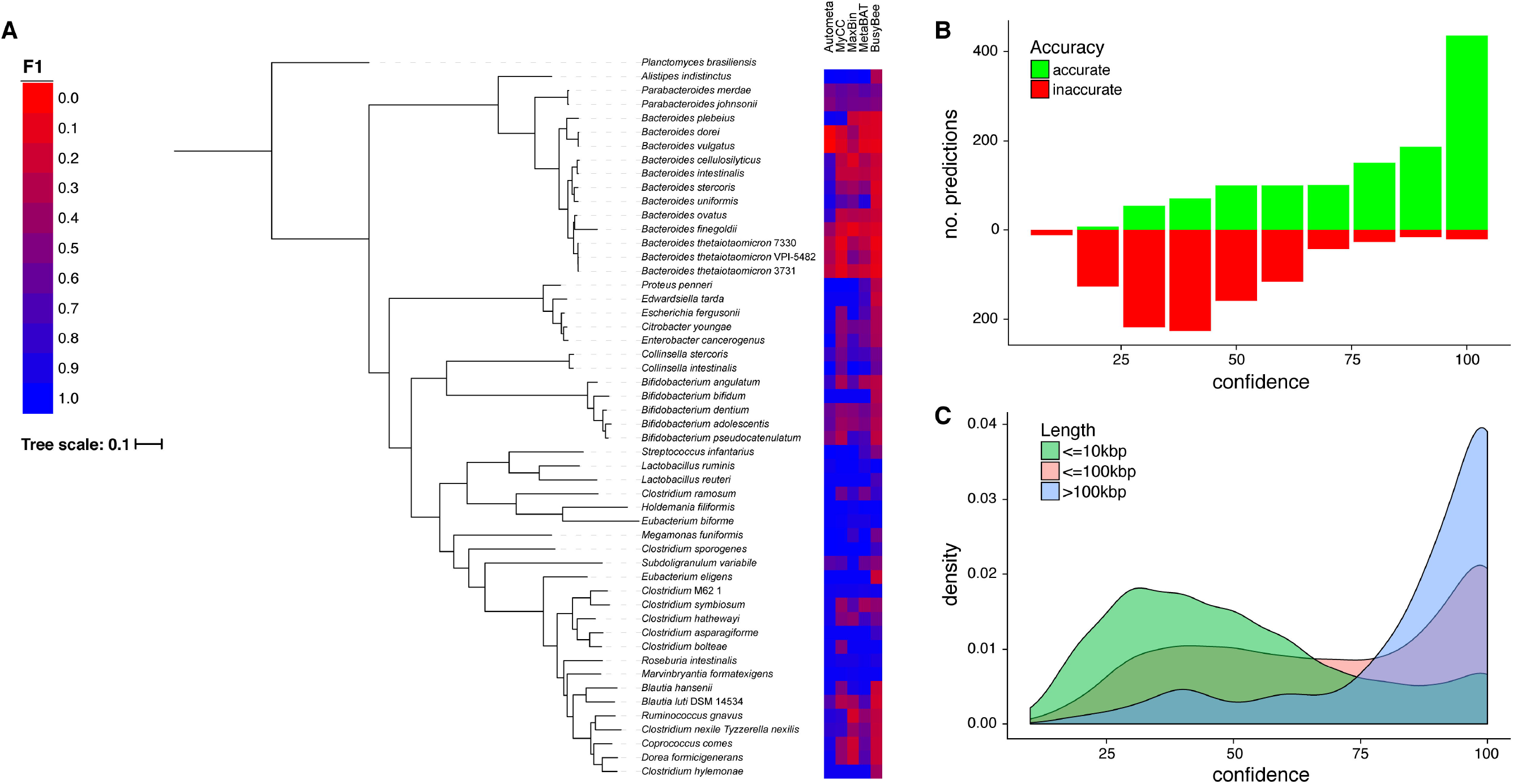
Performance testing and machine learning classification proof of concept using a synthetic metagenome with high coverage and strain overlap. (A) F1 values of individual genomes in Mix-51 as compared to phylogeny based on concatenated protein marker alignments using AMPHORA2 (Wu and Scott, 2012). (B) Number of accurate and inaccurate predictions compared to the confidence (based on jackknife cross validation, **Figure S13**) of the decision tree classifier when the known reference genomes of single copy marker-containing contigs are provided as a ground truth. In other words, when the model is provided accurate assignments of the marker-containing contigs, its confidence is well correlated to its accuracy (Pearson Correlation Coefficient for the percent of accurate predictions is 0.9887931, *p* = 6.809 × 10^−8^). (C) Density plot showing confidence of the classifier’s predictions compared to the length of the contig being classified.

In addition to stress testing these automated binning programs with high-strain overlap, the Mix-51 sample was used to validate the performance of the machine learning classification step as a proof of concept. When contigs containing single copy marker contigs (Rinke *et al.*, 2013) were used to train the decision tree classifier (with known reference genomes--as annotated by metaquast alignment (Mikheenko *et al.*, 2016)--provided as labels), the classifier was able to predict the genome identity of other contigs with very high accuracy, where predictions were reported to have high confidence values. There was a strong correlation between the classifier’s confidence (as determined by a Jackknife Cross validation approach (**Figure S13**) and the percent of accurate predictions (**Figure 3A**; Pearson Correlation Coefficient, 0.9887931, *p* = 6.809 × 10^−8^). For instance, 95.4% (436/457) of predictions with 100% confidence were accurate, recruiting 17 Mbp of sequence. On the other hand, only 18.3% (131/714) of predictions with less than 50% confidence were accurate. It also appears that the confidence of the classifier is positively associated with sequence length (**Figure 3B**), likely because the signal and resolving power of *k*-mer frequency is known to improve with sequence length (Sangwan *et al.*, 2016). The median confidence of predictions for contigs greater than 100 kbp, 10 - 100 kbp, and less than 10 kbp were 90%, 70%, and 50%, respectively (**Figure 3B**).

### 3.3 Performance in uniform coverage simulated metagenomes

Due to the innate complexity of some marine invertebrate associated microbial communities, such as marine sponges (Hentschel *et al.*, 2012; Taylor *et al.*, 2007; Lackner *et al.*, 2017), we tested the scalability of composition and homology-based techniques for single sample binning analysis. To this end we tested our algorithm along with MyCC, MaxBin, MetaBAT, and BusyBee Web using a set of increasingly complex simulated sequence sets with uniform coverage (**Table 1**).

For each of the simulated datasets, Autometa was able to recover more genome bins (**Table S8**), including in the largest tested dataset (10,000 Mbp), which represented the simulated sequencing of a metagenome containing 2,617 bacterial genomes. At the same time, the median F1 score of bins yielded by Autometa is consistently close to 1.0 (**Table S9**), up to and including the 5,000 Mbp dataset (**Figure 4A**). It should be noted that MaxBin was the only pipeline other than Autometa able to complete successfully for the two largest datasets (5,000 Mbp and 10,000 Mbp). In the smaller datasets (78.125 Mbp, 156.25 Mbp and 312.5 Mbp), the performance of other pipelines is comparable to Autometa, but their performance rapidly declines in more complex datasets. We also calculated F1 recovery for all pipelines (the sum of all F1 scores for each genome bin divided by the theoretical maximum sum; **Figure 4B, Table 1**). Based on this metric, Autometa and MyCC performed comparably for the three smallest datasets. However, for the larger datasets (625 Mbp to 10,000 Mbp) Autometa consistently outperformed MyCC. MaxBin and MetaBAT underperformed Autometa and MyCC in all datasets except for the smallest 78.125 Mbp by this measure (**Table S10**).

**Figure 4.**
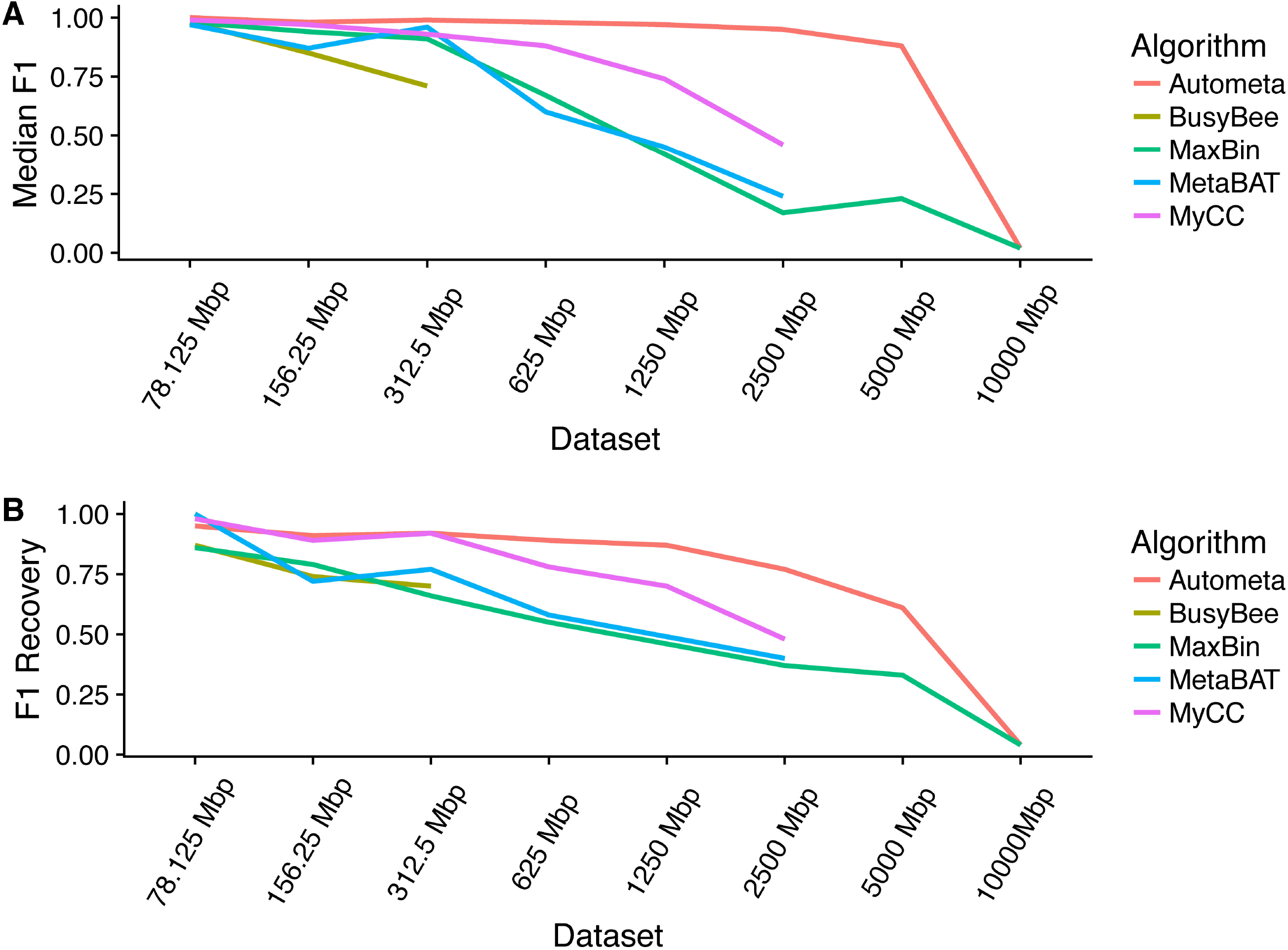
Performance in increasingly complex simulated metagenomes with uniform sequence coverage profiles based on median F1 and F1 recovery.

It is worth noting that while Autometa’s performance appears to drop dramatically after the 5000 Mbp dataset, this drop is most likely a result of the sharp decline in the assembly quality (**Table 1**), whereby the N_50_ (for contigs ≥3 kbp) drops from 7,222 to 4,611 bp, and where the total assembled length drops from 3179.1 Mbp to 427.5 Mbp (for contigs ≥3 kbp), for the 5,000 Mbp and 10,000 Mbp datasets, respectively. It is possible that if simulated sequencing parameters were adjusted to simulate greater sequencing depth, the quality of the assembly and thus binning results would continue to scale.

## 4 Discussion

Shotgun sequencing coupled with genome binning enables species-level resolution of metagenomes, even when the genomes of their microbial constituents lack representatives in reference databases. This reference-free approach is moving the field of microbiology from phylogenetic profiling of communities and aggregate interpretation of metagenomic data to a higher resolution perspective of which organisms play what role in a given ecosystem. Such information can be invaluable in a diverse array of biotechnological applications, such as identifying the source of bioactive secondary metabolites in complex marine invertebrate communities (Piel *et al.*, 2004; Miller *et al.*, 2017; Lopera *et al.*, 2017), or antibiotic resistance mechanisms (Ashton *et al.*, 2015; Shore *et al.*, 2011; Moss *et al.*, 2017; Loman *et al.*, 2013; van der Helm *et al.*, 2017) in uncultured clinical samples (Grumaz et al., 2016; Doughty *et al.*, 2014). However, despite the advances stemming from this paradigm shift in metagenomic analysis, a number of challenges remain.

Many available automated binning programs require the use of multiple samples in order to bin contigs into genome bins based on differential coverage profiles. However, this type of sample collection strategy is often not possible for marine invertebrate communities with dynamic compositions (Miller, Weyna, *et al.*, 2016), and can be too costly for exploratory sequencing studies (Miller *et al.*, 2017). Furthermore, these techniques typically rely on the use of co-assemblies, a strategy whereby reads from multiple samples are pooled prior to assembly. This approach leads to increasingly aggregate and chimeric representation of sequences and has been shown to reduce the overall genome assembly quality of constituent genomes (Olm *et al.*, 2017), which, in turn, reduces the accuracy of the genome binning process.

In a recent review by Sangwan *et al.*, the authors cited a general lack of binning strategies that integrate phylogenetic analysis with nucleotide composition (Sangwan *et al.*, 2016). Part of the challenge in analyzing marine invertebrate-associated microbiomes is the fact that most marine invertebrates lack any type of reference genome, and thus, unlike studies of the human microbiome, resulting reads from shotgun sequencing cannot be easily separated from these sequencing datasets with alignment techniques. Much of our approach in developing Autometa aimed to both address this fundamental issue and to further leverage contig-level taxonomic assignments to improve the binning process. Other efforts have focused on removing prokaryotic contamination following *de novo* assembly efforts of eukaryotic genomes (Fierst and Murdock, 2017), while here we have implemented kingdom-level taxonomic partitioning prior to binning, in addition to incorporating taxonomic information in clustering and classification steps, improved binning performance both for Autometa and for other tested binning pipelines in the bryozoan metagenome tested here.

The necessity for genome binning ultimately stems from the underlying shortcomings of modern sequencing technology (Sangwan *et al.*, 2016; Miller *et al.*, 2017), especially in regards to the trade off between read length, accuracy, and sequencing depth (Loman and Pallen, 2015). Thus, short read sequencing technologies, such as Illumina, are the only platforms currently capable of delivering sufficient sequencing depth and per-read accuracy to effectively assemble low abundance genomes directly from host-associated metagenomes without physical or chemical enrichment of bacterial DNA (Quince *et al.*, 2017), which can introduce unforseen sampling bias. As the throughput and accuracy of longer read technology continue to advance, the demands for advanced binning strategies could feasibly decrease. However, for the foreseeable future, genome-resolved metagenomics will rely on contig-binning strategies based on a combination of coverage, composition, and homology. No individual binning model will likely be able to outperform all others under every circumstance. Furthermore, there are fundamental limitations of binning sequences based on coverage, composition, and homology features that complicate the proper assignment of mobile elements such as plasmids and genes acquired through horizontal transmission, especially when they are poorly assembled (for instance, due to high repeat content). Thus, it is important that users understand the assumptions of each approach (Sangwan *et al.*, 2016; Sedlar *et al.*, 2017) and interpret results accordingly. Others have suggested and demonstrated that a combined strategy, particularly using programs with distinct underlying algorithms, is most likely to yield the most robust results (Sangwan *et al.*, 2016; C. M. K. Sieber *et al.*, 2017; Song and Thomas, 2017). We have shown here, however, that the integration of taxonomic information with nucleotide composition in Autometa allows it to outperform several other pipelines in host-associated and extremely complex metagenomes, yielding hundreds of high-quality genome bins from single datasets. This capability will complement existing multi-sample techniques by allowing the analysis of inter-sample strain variability in high resolution, which is likely to be seen in vertically-transmitted symbionts.

## 5 Acknowledgements

Work in J.C.K.’s lab is supported by R21AI121704 from NIAID as well as the School of Pharmacy, the Graduate School, and the Institute for Clinical & Translational Research at the University of Wisconsin-Madison. I.J.M. is supported by an American Foundation for Pharmaceutical Education Predoctoral Fellowship and E.R.R. is supported by the Biotechnology Training Program, supported by NIGMS, award number T32GM008349. This research was performed in part using the computer resources and assistance of the UW-Madison Center For High Throughput Computing (CHTC) in the Department of Computer Sciences. The CHTC is supported by UW-Madison, the Advanced Computing Initiative, the Wisconsin Alumni Research Foundation, the Wisconsin Institutes for Discovery, and the National Science Foundation, and is an active member of the Open Science Grid, which is supported by the National Science Foundation and the U.S. Department of Energy’s Office of Science. The authors wish to thank Miguel Pignatelli for insight into his Blast2LCA algorithm. The authors would also like thank Marc Chevrette, Chase Clark and Cedric Laczny for helpful feedback.

## References

Albertsen, M. et al. (2013) Genome sequences of rare, uncultured bacteria obtained by differential coverage binning of multiple metagenomes. Nat. Biotechnol., 31, 533–538.

Alivisatos, A.P. et al. (2015) A unified initiative to harness Earth’s microbiomes. Science, 350, 507–508.

Alneberg, J. et al. (2014) Binning metagenomic contigs by coverage and composition. Nat. Methods, 11, 1144–1146.

Ashton, P.M. et al. (2015) MinlON nanopore sequencing identifies the position and structure of a bacterial antibiotic resistance island. Nat. Biotechnol., 33, 296–300.

Bennett, G.M. and Moran, N.A. (2015) Heritable symbiosis: The advantages and perils of an evolutionary rabbit hole. Proc. Natl. Acad. Sci. U. S. A., 112, 10169–10176.

Bolger, A.M. et al. (2014) Trimmomatic: A flexible trimmer for Illumina sequence data. Bioinformatics, 30, 2114–2120.

Brown, C.T. et al. (2015) Unusual biology across a group comprising more than 15% of domain Bacteria. Nature, 523, 208–211.

Buchfink, B. et al. (2015) Fast and sensitive protein alignment using DIAMOND. Nat. Methods, 12, 59–60.

Buick, R. (2008) When did oxygenic photosynthesis evolve? Philos. Trans. R. Soc. Lond. B Biol. Sci., 363, 2731–2743.

Cao, Y. and Wang, L. (2017) Automatic selection of t-SNE Perplexity. arXiv:1708.03229 [cs.AI].

Chevrette, M.G. et al. (2017) SANDPUMA: Ensemble predictions of nonribosomal peptide chemistry reveals biosynthetic diversity across *Actinobacteria*. Bioinformatics, 33, 3202–3210.

Dick, G.J. et al. (2009) Community-wide analysis of microbial genome sequence signatures. Genome Biol., 10, R85.

Doughty, E.L. et al. (2014) Culture-independent detection and characterisation of *Mycobacterium tuberculosis* and *M. africanum* in sputum samples using shotgun metagenomics on a benchtop sequencer. PeerJ, 2, e585.

Dubilier, N. et al. (2015) Microbiology: Create a global microbiome effort. Nature, 526, 631–634.

Escobar-Zepeda, A. et al. (2015) The road to metagenomics: From microbiology to DNA sequencing technologies and bioinformatics. Front. Genet., 6, 348.

Ester, M. et al. (1996) A density-based algorithm for discovering clusters in large spatial databases with noise. In, Simoudis, E. et al. (eds), Proceedings of the second international conference on knowledge discovery and data mining., pp. 226–231.

Fierst, J.L. and Murdock, D.A. (2017) Decontaminating eukaryotic genome assemblies with machine learning. BMC Bioinformatics, 18, 533.

Grumaz, S. et al. (2016) Next-generation sequencing diagnostics of bacteremia in septic patients. Genome Med., 8, 73.

Heintz-Buschart, A. et al. (2016) Integrated multi-omics of the human gut microbiome in a case study of familial type 1 diabetes. Nat Microbiol, 2, 16180.

van der Helm, E. et al. (2017) Rapid resistome mapping using nanopore sequencing. Nucleic Acids Res., 45, e61.

Hentschel, U. et al. (2012) Genomic insights into the marine sponge microbiome. Nat. Rev. Microbiol., 10, 641–654.

Huang, W. et al. (2012) ART: A next-generation sequencing read simulator. Bioinformatics, 28, 593–594.

Hyatt, D. et al. (2010) Prodigal: Prokaryotic gene recognition and translation initiation site identification. BMC Bioinformatics, 11, 119.

Kang, D.D. et al. (2015) MetaBAT, an efficient tool for accurately reconstructing single genomes from complex microbial communities. PeerJ, 3, e1165.

Lackner, G. et al. (2017) Insights into the lifestyle of uncultured bacterial natural product factories associated with marine sponges. Proc. Natl. Acad. Sci. U. S. A., 114, e347–356.

Laczny, C.C. et al. (2014) Alignment-free visualization of metagenomic data by nonlinear dimension reduction. Sci. Rep., 4, 4516.

Laczny, C.C. et al. (2017) BusyBee Web: Metagenomic data analysis by bootstrapped supervised binning and annotation. Nucleic Acids Res., 45, e171–179.

Laczny, C.C. et al. (2015) VizBin - An application for reference-independent visualization and human-augmented binning of metagenomic data. Microbiome, 3, 1.

Lin, H.-H. and Liao, Y.-C. (2016) Accurate binning of metagenomic contigs via automated clustering sequences using information of genomic signatures and marker genes. Sci. Rep., 6, 24175.

Liu, Y. et al. (2010) Understanding of internal clustering validation measures. In, 2010 IEEE International Conference on Data Mining., pp. 911–916.

Loman, N.J. et al. (2013) A culture-independent sequence-based metagenomics approach to the investigation of an outbreak of Shiga-toxigenic *Escherichia coli* O104: H4. JAMA, 309, 1502–1510.

Loman, N.J. and Pallen, M.J. (2015) Twenty years of bacterial genome sequencing. Nat. Rev. Microbiol., 13, 787–794.

Lopera, J. et al. (2017) Increased biosynthetic gene dosage in a genome-reduced defensive bacterial symbiont. mSystems, 2, e00096–17.

McCutcheon, J.P. and Moran, N.A. (2012) Extreme genome reduction in symbiotic bacteria. Nat. Rev. Microbiol., 10, 13–26.

McNulty, N.P. et al. (2011) The impact of a consortium of fermented milk strains on the gut microbiome of gnotobiotic mice and monozygotic twins. Sci. Transl. Med., 3, 106ra106.

Medini, D. et al. (2005) The microbial pan-genome. Curr. Opin. Genet. Dev., 15, 589–594.

Mikheenko, A. et al. (2016) MetaQUAST: Evaluation of metagenome assemblies. Bioinformatics, 32, 1088–1090.

Miller, I.J. et al. (2017) Interpreting microbial biosynthesis in the genomic age: Biological and practical considerations. Mar. Drugs, 15, 165.

Miller, I.J., Vanee, N., et al. (2016) Lack of overt genome reduction in the bryostatin-producing bryozoan symbiont *‘Candidatus* Endobugula sertula’. Appl. Environ. Microbiol., 82, 6573–6583.

Miller, I.J., Weyna, T.R., et al. (2016) Single sample resolution of rare microbial dark matter in a marine invertebrate metagenome. Sci. Rep., 6, 34362.

Moss, E. et al. (2017) *De novo* assembly of microbial genomes from human gut metagenomes using barcoded short read sequences. bioRxiv, 125211. doi: 10.1101/125211

Nielsen, H.B. et al. (2014) Identification and assembly of genomes and genetic elements in complex metagenomic samples without using reference genomes. Nat. Biotechnol., 32, 822–828.

Nurk, S. et al. (2017) metaSPAdes: A new versatile metagenomic assembler. Genome Res., 27, 824–834.

Olm, M.R. et al. (2017) dRep: A tool for fast and accurate genomic comparisons that enables improved genome recovery from metagenomes through de-replication. ISME J., 11, 2864–2868.

Parks, D.H. et al. (2015) CheckM: Assessing the quality of microbial genomes recovered from isolates, single cells, and metagenomes. Genome Res., 25, 1043–1055.

Piel, J. et al. (2004) Antitumor polyketide biosynthesis by an uncultivated bacterial symbiont of the marine sponge *Theonella swinhoei*. Proc. Natl. Acad. Sci. U. S. A., 101, 16222–16227.

Quince, C. et al. (2017) Shotgun metagenomics, from sampling to analysis. Nat. Biotechnol., 35, 833–844.

Rinke, C. et al. (2013) Insights into the phylogeny and coding potential of microbial dark matter. Nature, 499, 431–437.

Sangwan, N. et al. (2016) Recovering complete and draft population genomes from metagenome datasets. Microbiome, 4, 8.

Sedlar, K. et al. (2017) Bioinformatics strategies for taxonomy independent binning and visualization of sequences in shotgun metagenomics. Comput. Struct. Biotechnol. J., 15, 48–55.

Shore, A.C. et al. (2011) Detection of staphylococcal cassette chromosome mec type XI carrying highly divergent mecA, mecl, mecR1, blaZ, and ccr genes in human clinical isolates of clonal complex 130 methicillin-resistant Staphylococcus aureus. Antimicrob. Agents Chemother., 55, 3765–3773.

Sieber, C.M.K. et al. (2017) Recovery of genomes from metagenomes via a dereplication, aggregation, and scoring strategy. bioRxiv, 107789. doi: 10.1101/107789

Sieber, K.B. et al. (2017) Lateral gene transfer between prokaryotes and eukaryotes. Exp. Cell Res., 358, 421–426.

Song, W.-Z. and Thomas, T. (2017) Binning_refiner: Improving genome bins through the combination of different binning programs. Bioinformatics, 33, 1873–1875.

Staley, J.T. and Konopka, A. (1985) Measurement of *in situ* activities of nonphotosynthetic microorganisms in aquatic and terrestrial habitats. Annu. Rev. Microbiol., 39, 321–346.

Taylor, M.W. et al. (2007) Sponge-associated microorganisms: Evolution, ecology, and biotechnological potential. Microbiol. Mol. Biol. Rev., 71, 295–347.

Teeling, H. et al. (2004) Application of tetranucleotide frequencies for the assignment of genomic fragments. Environ. Microbiol., 6, 938–947.

Tettelin, H. et al. (2008) Comparative genomics: The bacterial pan-genome. Curr. Opin. Microbiol., 11, 472–477.

Van Der Maaten, L. (2014) Accelerating t-SNE using tree-based algorithms. J. Mach. Learn. Res., 15, 3221–3245.

Wu, M. and Scott, A.J. (2012) Phylogenomic analysis of bacterial and archaeal sequences with AMPHORA2. Bioinformatics, 28, 1033–1034.

Wu, Y.-W. et al. (2014) MaxBin: An automated binning method to recover individual genomes from metagenomes using an expectation-maximization algorithm. Microbiome, 2, 26.

